# In-silico testing of new pharmacology for restoring inhibition and human cortical function in depression

**DOI:** 10.1101/2023.02.22.529541

**Authors:** Alexandre Guet-McCreight, Homeira Moradi Chameh, Frank Mazza, Thomas D. Prevot, Taufik A. Valiante, Etienne Sibille, Etay Hay

## Abstract

Reduced inhibition by somatostatin-expressing interneurons is associated with depression. Administration of positive allosteric modulators of α5 subunit-containing GABA_A_ receptor (α5-PAM) that selectively target this lost inhibition exhibit antidepressant and pro-cognitive effects in rodent models of chronic stress. However, the functional effects of α5-PAM on the human brain *in vivo* are unknown, and currently cannot be assessed experimentally. We modeled the effects of α5-PAM on tonic inhibition as measured in human neurons, and tested *in silico* α5-PAM effects on detailed models of human cortical microcircuits in health and depression. We found that α5-PAM effectively recovered impaired cortical processing as quantified by stimulus detection metrics, and also recovered the power spectral density profile of the microcircuit EEG signals. We performed an α5-PAM dose response and identified simulated EEG biomarker candidates. Our results serve to de-risk and facilitate α5-PAM translation and provide biomarkers in non-invasive brain signals for monitoring target engagement and drug efficacy.

## Introduction

A loss of cortical inhibition is associated with major depressive disorder (depression) (Levinson et al. 2010), and studies indicate the involvement of somatostatin-expressing (SST) inhibitory interneurons (Duman, Sanacora, and Krystal 2019; Fee et al. 2021; Fee, Banasr, and Sibille 2017; Fuchs et al. 2017; Lin and Sibille 2013, 2015; Northoff and Sibille 2014; Prevot and Sibille 2021; Seney et al. 2015; Song, Yoon, and Lee 2021). Functionally, cortical SST interneurons mediate lateral inhibition through inhibitory disynaptic loops (Obermayer et al. 2018; Silberberg and Markram 2007) and provide a “blanket of inhibition” that maintains low Pyr neuron spike rates at baseline (Gentet et al. 2012; Karnani et al. 2016; Karnani, Agetsuma, and Yuste 2014). SST interneurons primarily target the apical dendrites of pyramidal (Pyr) neurons, where they provide both synaptic and extrasynaptic (i.e., tonic) inhibition via the α5 subunit of GABA_A_ (α5-GABA_A_) receptors (Ali and Thomson 2008; Davenport et al. 2021; Donato et al. 2023; Schulz et al. 2018). While mostly studied in rodents, studies showed that α5-GABA_A_ is similarly expressed in human cortical pyramidal neurons and negligibly in interneurons (Hu et al. 2018). Accordingly, tonic inhibitory currents generated by α5-GABA_A_ receptors have been recorded in human cortical pyramidal neurons and are mostly absent in interneurons (Scimemi et al. 2006).

Novel benzodiazepine-like compounds with preferential affinities and positive allosteric modulation of α5-GABA_A_ receptors (α5-PAM) have been shown to elicit anxiolytic, antidepressant, and pro-cognitive effects in rodent models associated with reduced dendritic inhibition, such as chronic stress or aging, and are therefore promising new treatments for depression (Bernardo et al. 2022; Gill et al. 2011; Jacob 2019; Koh, Rosenzweig-Lipson, and Gallagher 2013; Piantadosi et al. 2016; Prevot et al. 2019). Comparatively, non-selective benzodiazepines (i.e., targeting a broader range of GABA_A_ receptor subunits; Sigel and Ernst, 2018) that are efficacious at treating anxiety (Gomez, Barthel, and Hofmann 2018) have several undesirable side-effects that include sedation, ataxia, amnesia, and abuse liability due to their broad modulation of cortical inhibition via α1 subunit-containing GABA_A_ receptors (McKernan et al. 2000; Rudolph and Möhler 2006), which are expressed more ubiquitously across neuron types (Hörtnagl et al. 2013; Nutt 2006). While α5-PAM effects on rodents suggest a potential role in the treatment of depression, their effects on human brain microcircuits remain unknown due to experimental and ethical limitations, thus meriting *in silico* testing.

There are two main factors that make testing α5-PAM effects on human microcircuits not trivial. First, although there are many similarities between cortical microcircuits in humans and other species, there are also important differences. Compared to rodents, human cortical circuits exhibit stronger excitatory and inhibitory synaptic connections (Campagnola et al. 2022; Komlósi et al. 2012; Molnár et al. 2016; Obermayer et al. 2018; Seeman et al. 2018), as well as larger morphological sizes and enhanced dendritic compartmentalization (Beaulieu-Laroche et al. 2018, 2021; Eyal et al. 2016; Gidon et al. 2020; Kalmbach et al. 2018, 2021). The effects of α5-PAM in rodents are therefore expected to translate to humans, but the extent to which species differences modify these effects remains to be determined. Another non-trivial factor in testing α5-PAM effects is the difference between the α5-PAM mode of operation and the underlying SST interneuron mechanisms of depression. α5-PAM in cortex acts specifically to enhance inhibition onto Pyr neuron apical dendrites (Ali and Thomson 2008; Davenport et al. 2021; Donato et al. 2023; Schulz et al. 2018), whereas reduced SST interneuron inhibition affects additional microcircuit connections from SST interneurons onto other interneuron subtypes (Donato et al. 2023; Yao et al. 2022). Thus, it is unclear whether α5-PAM is sufficient to fully recover circuit activity dynamics following reduced SST interneuron inhibition in depression.

Here we test *in silico* the effects of α5-PAM on human microcircuit spiking, function in terms of signal detection, and EEG signals, using our previous detailed models of human cortical microcircuits in health and depression (Yao et al., 2022). We characterize the effects of different doses of α5-PAM, identify biomarker candidates in EEG signals and establish their specificity compared to non-selective GABA_A_ receptor PAMs.

## Results

### Data-driven models of α5-PAM modulation in human neurons and in-silico testing of microcircuit function recovery

We first experimentally recorded tonic inhibition currents in single healthy human pyramidal neurons in the presence of GABA only, and in the presence of α5-PAM + GABA, and found a 52% increase in current magnitude during application of a reference dose of α5-PAM (3 µM) in addition to GABA (-65.9 ± 30.8 pA vs -95.3 ± 43.6 pA, paired-sample t-test, *p* < 0.05, Cohen’s *d* = 0.82; **Fig. 1A,C**). We then modeled the α5-PAM modulation in single neurons by constraining our previous human L2/3 Pyr neuron model to reproduce the average currents recorded *in-vitro* (**Fig. 1B-C**). We first fitted the tonic inhibition conductance (*G_tonic_*) in the neuron (uniformly in soma, basal dendrites, and apical dendrites) to reproduce the current magnitude recorded when applying GABA, and then fitted the α5-PAM modulation of *G_tonic_* in the apical dendrites to reproduce the 52% increase in tonic current magnitude. We thus estimated the α5-PAM modulation to be a 60% increase in conductance (**Fig. 1D**).

**Figure 1.**
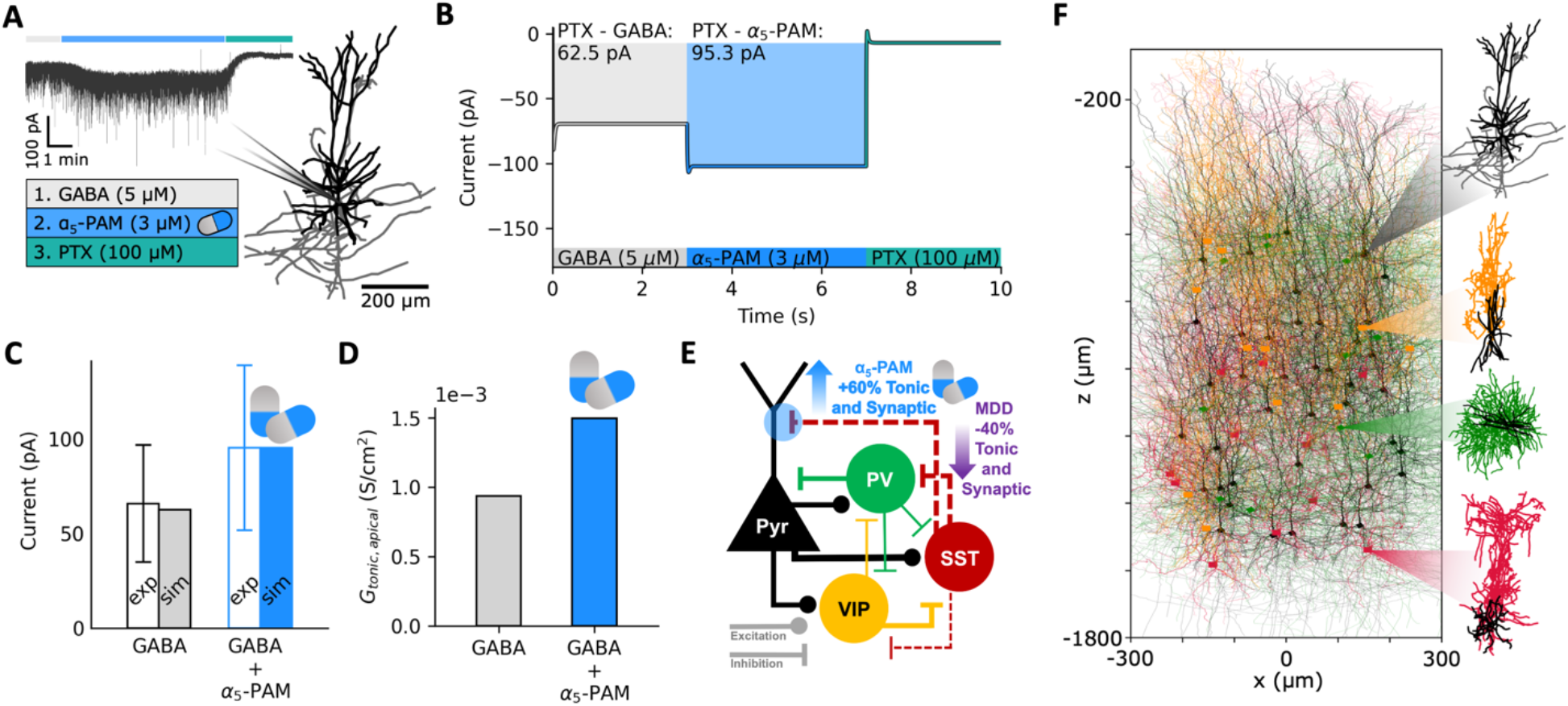
Data-driven simulation of α5-PAM effect on human neurons and microcircuits. **A.** Illustration of the detailed human L2/3 pyr neuron model used in the study and an example voltage-clamp recording under the different experimental conditions (GABA: application of GABA, α5-PAM: application of α5-PAM + GABA, PTX: GABA block). **B.** Simulated tonic inhibition current recordings at soma in the different conditions, fitted to reproduce the experimentally recorded current magnitude averages. **C.** Comparison of experimental and simulated current output magnitudes (relative to the output magnitude in the PTX condition) in the GABA (gray) and α5-PAM + GABA (blue) conditions. **D.** Derived apical tonic inhibition conductance values in the GABA (gray) and α5-PAM + GABA (blue) conditions. **E**. Schematic of the model L2/3 microcircuit connectivity and summary of depression (MDD) and derived α5-PAM effects on the microcircuit. **F**. Illustration of the detailed L2/3 microcircuit models with human model morphologies (from top to bottom: Pyr, VIP, PV, SST, color-coded as in E).

We next integrated this modulation into our previous biophysically detailed models of human L2/3 microcircuits in health and depression (Yao et al. 2022). The microcircuit models included key neuron types (Pyr, PV, SST, and VIP neurons), and the depression microcircuits involved a 40% reduction in SST interneuron synaptic and tonic inhibition onto the other neurons (**Fig. 1E-F**). We also applied the estimated α5-PAM modulation to synaptic inhibition mediated by SST interneurons onto Pyr neuron apical dendrites, since these connection types in cortex are α5-mediated.

We next tested the α5-PAM effects on the human cortical microcircuits by simulating microcircuit baseline and response to brief stimuli in health, depression, and depression + α5-PAM (**Fig. 2**). As we showed previously, compared to healthy circuits, reduced SST interneuron inhibition in depression microcircuits resulted in increased baseline spike rates (**Fig. 2A-B**; healthy: 0.75 ± 0.04 Hz; depression: 1.20 ± 0.06 Hz; paired-sample t-test, *p* < 0.05, Cohen’s *d* = 8.6), decreased SNR (**Fig. 2C**; healthy: 2.99 ± 0.73; depression: 2.00 ± 0.41; paired sample t-test, *p* < 0.05, Cohen’s *d* = -1.7), and worsened microcircuit function in terms of failed and false stimulus detection rates, calculated based on the distribution of Pyr neuron firing at baseline vs response averaged over 50 ms windows (**Fig. 2D-E**; healthy: 1.45 ± 0.47% and 1.54 ± 0.63%, depression: 5.71 ± 1.10% and 8.44 ± 2.53%, paired-sample t-test, *p* < 0.05, Cohen’s *d* = 5.0 and 3.7 respectively). Modulation of inhibition by a reference dose of α5-PAM in simulated depression microcircuits restored baseline spike rates (**Fig. 2B**; 0.76 ± 0.04 Hz, Cohen’s *d* = 0.2) and consequently SNR (**Fig. 2C**; 2.71 ± 0.63, Cohen’s *d* = -0.4) back to healthy levels. In both the depression and α5-PAM conditions, there were no effects on post-stimulus firing rate (**Fig. 2B**). The simulated reference dose of α5-PAM also recovered the failed and false stimulus detection rates nearly back to healthy level (**Fig. 2D-E;** 2.91 ± 0.83% and 2.59 ± 0.97%, Cohen’s *d* = 2.2 and 1.3).

**Figure 2.**
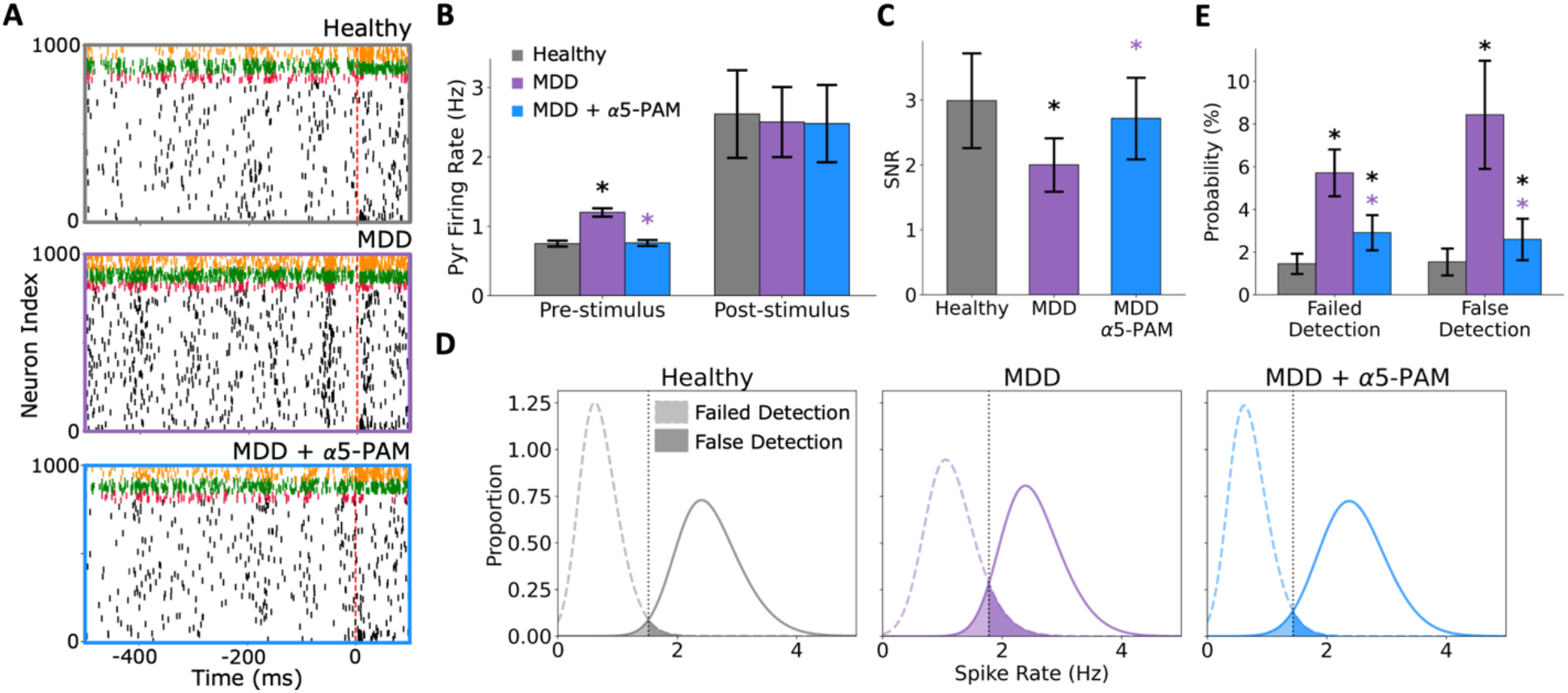
*In silico* application of α5-PAM in depression microcircuits restores healthy spike rates and function. **A.** Example raster plots of simulated baseline spiking and response to brief stimulus in healthy (top), depression (middle), and depression + α5-PAM (bottom) microcircuit models. Dashed line indicates stimulus time. Cell type color code is the same as in figure 1E-F. **B.** Pre- and post-stimulus Pyr neuron firing rates in the different simulated conditions. α5-PAM restores healthy levels of pre-stimulus firing. **C.** SNR of response in each condition, where α5-PAM boosts SNR to healthy levels. **D.** Distributions of pre- and post-stimulus firing rates. The vertical lines denote the decision boundaries, and the shaded areas show the failed/false detections. **E.** Probability of failed detection and false detection in each simulated condition. α5-PAM significantly reduced failed/false detection rates, bringing them close to the healthy levels. All asterisks denote significant paired t-tests (*p* < 0.05) with effect sizes greater than 1, when compared to healthy (black asterisks) or depression (MDD; purple asterisks). n = 200 randomized microcircuits per condition.

To test the effectiveness of higher or lower α5-PAM doses than the reference dose at recovering microcircuit function, we simulated α5-PAM modulation of apical inhibition ranging from 25-150% of the estimated modulation by the experimental reference dose (**Fig. 3**). The reference dose of α5-PAM (referred to as 100%) turned out to be the optimal dose at recovering baseline spike rates back to healthy levels in the simulated microcircuits (healthy: 0.75 ± 0.04 Hz; 100% α5-PAM: 0.76 ± 0.04 Hz, Cohen’s *d* = 0.2), whereas lower doses were not sufficient (25% α5-PAM: 1.07 ± 0.05 Hz, paired-sample t-test, *p* < 0.05, Cohen’s *d* = 6.5) and higher doses over-reduced the spike rate (150% α5-PAM: 0.60 ± 0.04 Hz, paired-sample t-test, *p* < 0.05, Cohen’s *d* = -3.8; **Fig. 3A**). The relationship between dose and effect on baseline rates was linear (Pearson correlation, r = -0.97, *p* < 0.05) and was only marginally better fitted by exponential or sigmoidal fits (∼10% improvement in the sum of squared errors). None of the doses tested had effects on post-stimulus firing rate (**Fig. 3A**). The relationship between dose and SNR was also linear (Pearson correlation, r = 0.49, *p* < 0.05; <5% difference in sum of squared errors between linear, sigmoid and exponential fits), with several doses restoring SNR back to healthy levels (100%-150%; **Fig. 3B**; healthy: 2.99 ± 0.73; 100% α5-PAM: 2.71 ± 0.63, Cohen’s *d* = -0.4; 125% α5-PAM: 2.82 ± 0.66, Cohen’s *d* = -0.2; 150% α5-PAM: 2.95 ± 0.74, Cohen’s *d* = -0.1). We note, however, that for doses greater than 100% the SNR was preserved because both baseline and response rates were similarly dampened, whereas the 100% dose effect involved minimal dampening and thus was more optimal. The relationship between dose and failed/false stimulus detection errors was nearly linear (Pearson correlation, r = -0.88 and -0.85, respectively, *p* < 0.05; < 10% difference in the sum of squared errors between linear, exponential and sigmoid fits), with a poor effect for low doses (25-50%; 25% α5-PAM: 6.36 ± 1.09% and 6.57 ± 1.95%, paired-sample t-test, *p* < 0.05, Cohen’s *d* = 5.8 and 3.5, compared to healthy), and errors rates recovering close to healthy levels for doses of 75-125% (**Fig. 3C**; 100% α5-PAM: 2.91 ± 0.83% and 2.59 ± 0.97%, paired-sample t-test, *p* < 0.05, Cohen’s *d* = 2.2 and 1.3, compared to healthy). The highest dose (150%) further reduced the failed and false error rates even below the healthy level (150% α5-PAM: 0.61 ± 0.32% and 0.73 ± 0.48%, paired-sample t-test, *p* < 0.05, Cohen’s *d* = -2.1 and - 1.4, compared to healthy), possibly through a greater dampening of pre-stimulus activity compared to post-stimulus when compared to healthy (Cohen’s *d* = -3.8 and -0.5, respectively).

**Figure 3.**
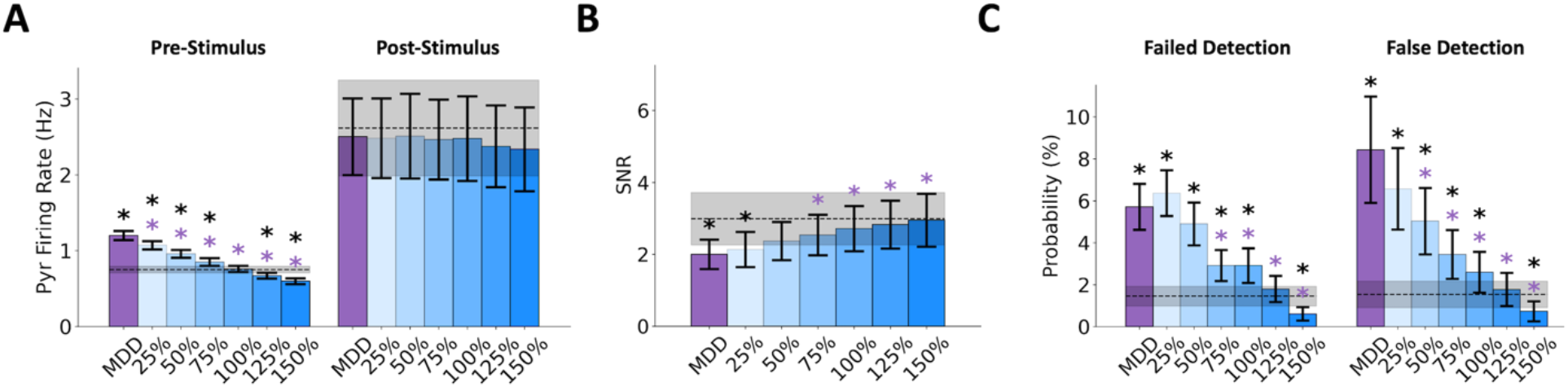
*In silico* dose-response highlights optimal levels for maximizing drug effects. **A-C.** Pyr neuron pre- and post-stimulus firing rates (A), SNR (B), and failed and false detection (C) for depression (MDD; purple) and different doses of α5-PAM (shades of blue: 25% - 150% of the estimated reference dose of α5-PAM). The black horizontal lines and shaded areas denote the healthy mean ± standard deviation. All asterisks denote significant paired t-tests (*p* < 0.05) with effect sizes greater than 1, when compared to healthy (black asterisks) or depression (MDD; purple asterisks). N = 200 randomized microcircuits per condition.

### In-silico EEG biomarkers of α5-PAM efficacy in depression microcircuits

We simulated EEG together with the microcircuit activity in health and depression conditions to identify signatures of α5-PAM efficacy in a clinically-relevant non-invasive signal (**Fig. 4A**). As we previously showed (Mazza et al. 2023), simulated EEG signals generated by the depression microcircuit models with reduced SST interneuron inhibition exhibited increased power in theta (healthy: 6.65 × 10^-14^ ± 6.34 × 10^-15^ mV^2^; depression: 9.34 × 10^-14^ ± 1.20 × 10^-14^ mV^2^; paired-sample t-test, *p* < 0.05, Cohen’s *d* = 2.8), alpha (healthy: 6.65 × 10^-14^ ± 7.32 × 10^-15^ mV^2^; depression: 9.12 × 10^-14^ ± 1.15 × 10^-14^ mV^2^; paired-sample t-test, *p* < 0.05, Cohen’s *d* = 2.5), and beta (Healthy: 3.84 × 10^-14^ ± 2.78 × 10^-15^ mV^2^; depression: 6.86 × 10^-14^ ± 4.99 × 10^-15^ mV^2^; paired-sample t-test, *p* < 0.05, Cohen’s *d* = 7.4) frequency bands (**Fig. 4B**). When α5-PAM was applied to the microcircuits at the reference dose, the power spectral density profile was restored close to the healthy at all frequency bands, except for a slight shift in theta band peak (**Fig. 4B**; 100% α5-PAM compared to healthy - θ: 7.41 × 10^-14^ ± 7.71 × 10^-15^ mV^2^, Cohen’s *d* = 1.1; α: 6.85 × 10^-14^ ± 8.44 × 10^-15^ mV^2^, Cohen’s *d* = 0.2; β: 4.12 × 10^-14^ ± 3.78 × 10^-15^ mV^2^, Cohen’s *d* = 0.8). There was a linear relationship between the effect of different α5-PAM doses at restoring power in theta (**Fig. 4C**; Pearson correlation, r = -0.67, *p* < 0.05; < 1% difference in the sum of squared errors between linear, exponential and sigmoid fits), alpha (**Fig. 4D**; Pearson correlation, r = -0.70, *p* < 0.05; < 1% difference in the sum of squared errors between linear, exponential and sigmoid fits), and beta (**Fig. 4E**; Pearson correlation, r = -0.92, *p* < 0.05; < 5% difference in the sum of squared errors between linear, exponential and sigmoid fits) frequency bands. The reference dose restored the power profile in alpha and beta bands (**Fig. 4D-E**), and a higher dose was required to restore the power profile in the theta band (**Fig. 4C**; 125% α5-PAM compared to healthy - θ: 6.90 × 10^-14^ ± 6.60 × 10^-15^ mV^2^, Cohen’s *d* = 0.4).

**Figure 4.**
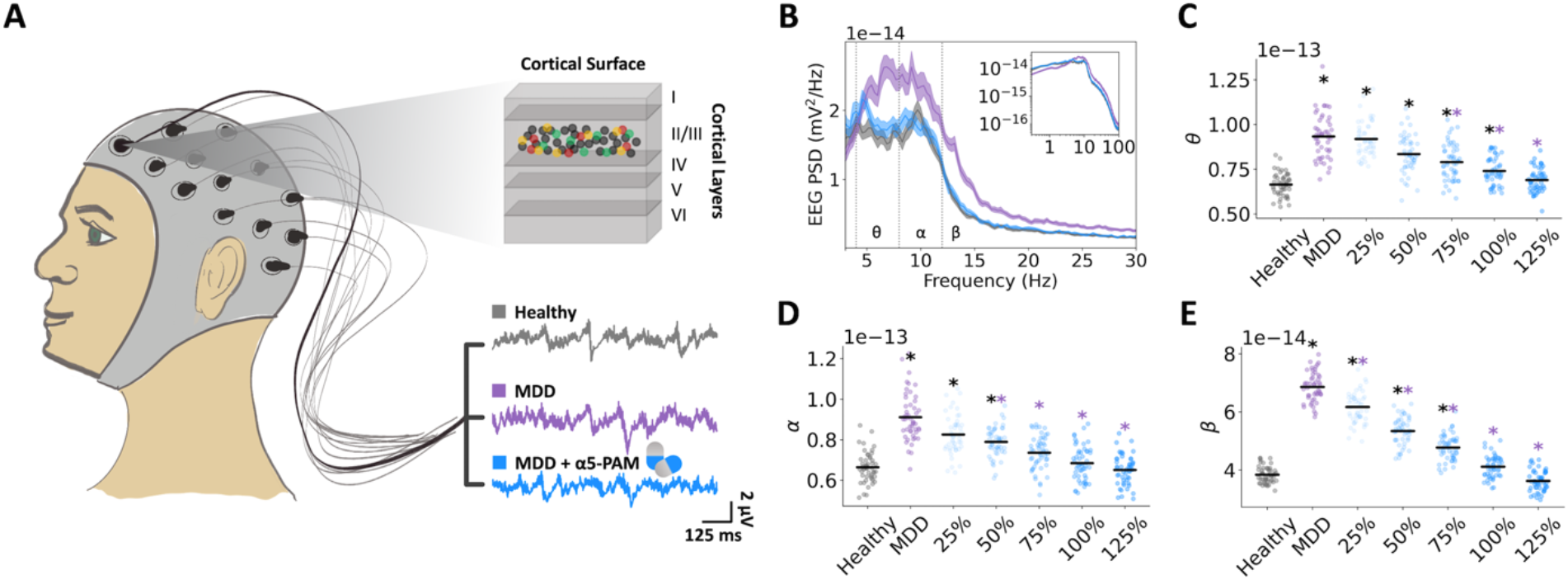
EEG power spectral biomarkers of α5-PAM efficacy. **A.** Illustration of EEG signals generated from the human cortical microcircuit models. **B.** Power spectral density (PSD) of simulated EEG from microcircuit models in each condition as labeled in A (bootstrapped mean, and 95% confidence intervals). α5-PAM restored the PSD profile to healthy level. Inset – PSD plotted in log scale. **C-E.** PSD in theta (4 - 8 Hz, C), alpha (8 - 12 Hz, D), and beta (12 - 21 Hz, E) bands for different α5-PAM doses (as in Fig. 3). All asterisks denote significant paired t-tests (*p* < 0.05) with effect sizes greater than 1, when compared to healthy (black asterisks) or depression (MDD; purple asterisks). N = 50 randomized microcircuits per condition.

To further assess features of the spectral biomarkers of α5-PAM efficacy, we decomposed the power spectral density profiles into aperiodic (**Fig. 5A-C**) and periodic (**Fig. 5D-F**) components. As we have shown previously (Mazza et al. 2023), depression microcircuits with reduced SST interneuron inhibition primarily exhibited altered aperiodic exponents (healthy: 0.96 ± 0.09 mV^2^/Hz; depression: 0.67 ± 0.07 mV^2^/Hz; paired-sample t-test, *p* < 0.05, Cohen’s *d* = -3.4) and broadband power (healthy: 1.03 × 10^-13^ ± 8.49 × 10^-15^ mV^2^; depression: 1.33 × 10^-13^ ± 1.04 × 10^-14^ mV^2^; paired-sample t-test, *p* < 0.05, Cohen’s *d* = 3.1) as well as increased periodic theta frequency power (healthy: 1.12 ± 0.18 mV^2^; depression: 1.53 ± 0.21 mV^2^; paired-sample t-test, *p* < 0.05, Cohen’s *d* = 2.0) and increased low-beta frequency power (healthy: 1.09 ± 0.28 mV^2^; depression: 1.73 ± 0.33 mV^2^; paired-sample t-test, *p* < 0.05, Cohen’s *d* = 2.0, **Fig. 5**). The reference α5-PAM dose (100%) restored the aperiodic power and exponent back to healthy level (100% α5-PAM, power: 1.11 × 10^-13^ ± 9.84 × 10^-15^ mV^2^, Cohen’s *d* = 0.8; exponent: 0.98 ± 0.09 mV^2^/Hz, Cohen’s *d* = 0.2 compared to healthy, **Fig. 5A-C**), as well as the periodic power in theta and alpha bands (100% α5-PAM, θ: 1.18 ± 0.26 mV^2^, Cohen’s *d* = 0.2; β: 1.10 ± 0.30 mV^2^, Cohen’s *d* = 0.03 compared to healthy, **Fig. 5D-F**). The relationship between α5-PAM dose and the effect on aperiodic or periodic components was linear (Pearson correlation, aperiodic power: r = -0.65, exponent: r = 0.71, θ: r = -0.48, β: r = -0.59; *p* < 0.05; < 6% difference in the sum of squared errors between linear, exponential, and sigmoid fits).

**Figure 5.**
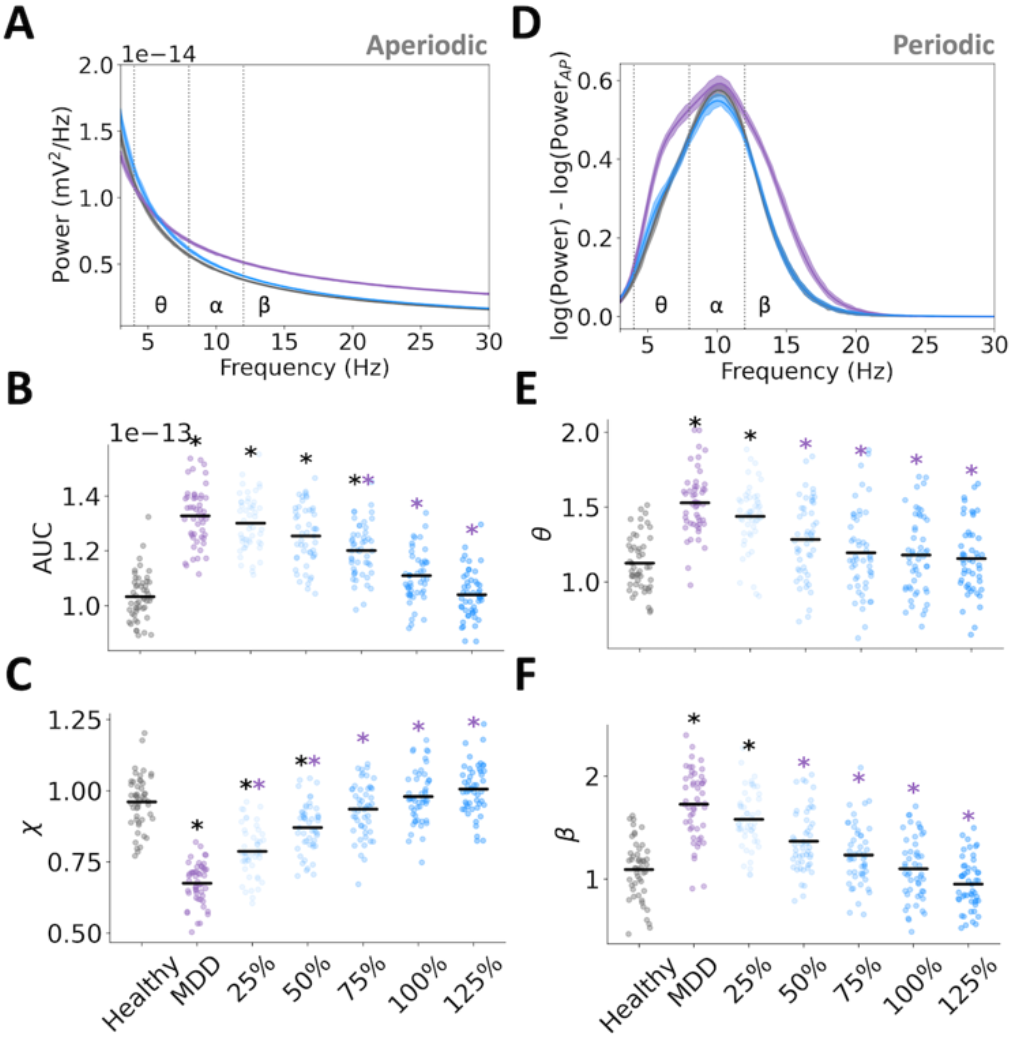
Decomposed EEG power spectral biomarkers of α5-PAM efficacy. **A.** Aperiodic components of the PSD for each condition (healthy: black; depression: purple; depression + 100% α5-PAM: blue). **B-C.** Broadband power spectral density area under the curve (AUC, 3-30 Hz, B) and exponent (χ, C) of the aperiodic component of the PSD for each α5-PAM dose (100%: fitted parameters for the 3 μM experimental α5-PAM dose). **D.** Periodic component of the PSD for each condition (healthy: black; depression: purple; depression + 100% α5-PAM: blue). **E-F.** Power of the periodic components of PSD in theta (θ, 4-8 Hz; E) and beta (β, 12-21 Hz; F) bands for each α5-PAM dose. All asterisks denote significant paired t-tests (*p* < 0.05) with effect sizes greater than 1, when compared to healthy (black asterisks) or depression (MDD; purple asterisks). N = 50 randomized microcircuits per condition.

We next compared the efficacy and EEG biomarkers of simulated α5-PAM with those of non-selective PAM, e.g. corresponding to a non-selective benzodiazepine (boosting a broad range of GABA_A_ subunits), by simulating a 60% increase in tonic and synaptic inhibition in all of the microcircuit inhibitory connections (**Fig. 6A**). Though simulated non-selective PAM reduced baseline microcircuit spike rates, it did not restore spike rates sufficiently back to healthy levels (**Fig. 6B**; non-selective PAM compared to healthy: 1.02 ± 0.06 Hz, paired-sample t-test, *p* < 0.05, Cohen’s *d* = 5.5). Accordingly, simulated non-selective PAM did not improve false detection rate (**Fig. 6C-D**; non-selective PAM: 6.78 ± 1.73%, healthy: 1.54 ± 0.63%, paired-sample t-test, *p* < 0.05, Cohen’s *d* = 4.0) and even worsened the failed detection rate (non-selective PAM: 7.99 ± 1.17%, healthy: 1.45 ± 0.47%, paired-sample t-test, *p* < 0.05, Cohen’s *d* = 7.3).

**Figure 6.**
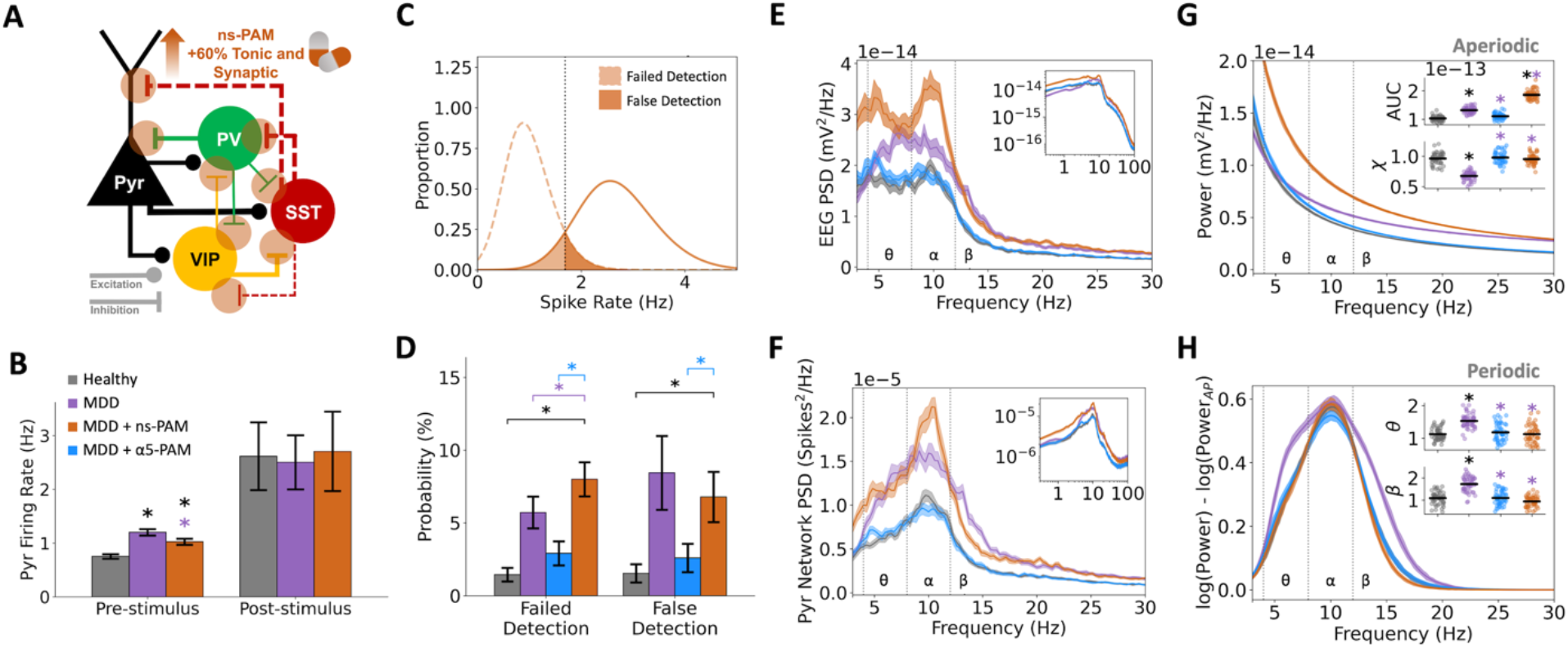
*In silico* application of non-selective PAM fails to recover microcircuit function and EEG. **A.** Schematic of non-selective PAM (ns-PAM) effects on the model microcircuit. **B.** Pre- and post-stimulus Pyr neuron firing rates in the different conditions. **C.** Distributions of pre- and post-stimulus firing rates. **D.** Probability of failed detection and false detection with non-selective PAM was worsened and unchanged, respectively, compared to the depression condition (MDD). **E.** EEG PSD, bootstrapped mean, and 95% confidence intervals. Inset: EEG PSD in log scale. **F.** Spikes PSD of Pyr neurons, bootstrapped mean, and 95% confidence intervals. Inset: spikes PSD in log scale**. G.** Fitted aperiodic components of the EEG PSD for each condition. Upper inset: broadband (3-30 Hz) area under the curve (AUC). Lower inset: exponent (χ). **H.** Fitted periodic component of the EEG PSD for each condition. Inset plots: integral of the power spectral density in the theta (θ) and beta (β) frequency ranges. All asterisks denote significant paired t-tests (*p* < 0.05) with effect sizes greater than 1, when compared to healthy (black asterisks) or depression (MDD; purple asterisks). n = 200 randomized microcircuits per condition for panels A-D. n = 50 randomized microcircuits per condition for panels E-H.

We analyzed the effects of non-selective PAM on simulated EEG power spectral density profile and found increased power in all frequency bands (**Fig. 6E**). Increased broadband power was similarly seen in the PSD of the Pyr neuron spiking (**Fig. 6F**). When decomposed into aperiodic and periodic components, we found that non-selective PAM caused a broadband upward shift in the aperiodic component (**Fig. 6G**, healthy: 1.03 × 10^-13^ ± 8.49 × 10^-15^ mV^2^; non-selective PAM: 1.86 × 10^-13^ ± 1.61 × 10^-14^ mV^2^; paired-sample t-test, *p* < 0.05, Cohen’s *d* = 6.4), but recovered the aperiodic exponent parameter (**Fig. 6G** inset, healthy: 0.96 ± 0.09 mV^2^/Hz; non-selective PAM: 0.95 ± 0.08 mV^2^/Hz; Cohen’s *d* = -0.1) as well as the periodic theta power (**Fig. 6H** inset, healthy: 1.12 ± 0.18 mV^2^; non-selective PAM: 1.12 ± 0.27 mV^2^; Cohen’s *d* = -0.01) and low-beta power (**Fig. 6H** inset, healthy: 1.09 ± 0.28 mV^2^; non-selective PAM: 0.94 ± 0.23 mV^2^; Cohen’s *d* = -0.6).

## Discussion

In this work, we tested *in silico* the effects of α5-PAM in detailed human cortical microcircuit models of depression and found that several indicators quantifying microcircuit dynamics, function, and EEG profile were brought back to healthy levels. We showed that the functional recovery, measured as failed and false detection, was on the same order of magnitude as the pro-cognitive effects measured in chronically stressed mice (Prevot et al. 2019). We further identified EEG biomarkers of different dose effects, highlighting recovery of EEG power in theta and beta frequencies, as well as broadband recovery. This mechanistic demonstration of α5-PAM efficacy on human cortical microcircuits could serve to guide pre-clinical studies, de-risk and facilitate translation to human clinical use, and provide candidate biomarkers in non-invasive brain signals for monitoring drug efficacy. In particular, our in-silico biomarker candidates can be tested in improving patient stratification and treatment outcome prediction, by identifying those that have the relevant depression EEG profile and that could benefit from being administered α5-PAM.

Our results demonstrated that α5-PAM could directly recover function and resting state EEG features associated with a loss of SST-mediated inhibition in depression (Mazza et al. 2023; Yao et al. 2022), despite only boosting SST→Pyr synapses and not SST→PV and SST→VIP synapses, which were also reduced in our depression models. This is likely because the loss of SST interneuron inhibition in depression has a direct and thus much larger impact on Pyr neurons than the indirect disinhibitory effect of reduced SST interneuron inhibition onto interneurons. In agreement with this rationale, we demonstrated that the use of a non-selective PAM failed to recover circuit activity to healthy levels, likely due to an indiscriminatory boosting of inhibition throughout the circuit rather than localized to the SST→Pyr connections. Similarly, while non-selective PAM dampened the elevated spike rates in depression to some extent, in line with previous studies during application of different non-selective benzodiazepines in rodent cortical cultures (Bader et al. 2017), the effects were small compared to α5-PAM effects, possibly as a result of the non-selective PAM boosting of all inhibitory connections in the microcircuit.

Recovery of theta, alpha, and lower beta frequency band power as a result of α5-PAM administration are relevant to depression diagnosis and severity, where these band powers are enhanced (de Aguiar Neto and Rosa 2019; Fernández-Palleiro et al. 2020; Grin-Yatsenko et al. 2010; Mazza et al. 2023; Newson and Thiagarajan 2019), and could be used as indicators of treatment response (Arns et al. 2015; Bailey et al. 2018; Bruder et al. 2008). We note that we did not observe beta peaks in our models, so the changes in the PSD lower beta band are likely more representative of a combination of changes in the alpha peak width and the aperiodic component. Our results demonstrate mechanistically that α5-PAM can directly recover resting state aperiodic and periodic power spectral biomarkers (i.e. corresponding to measures of asynchronous and synchronous microcircuit dynamics, respectively) associated with a loss of SST-mediated inhibition in models of depression (Mazza et al. 2023). Across all functional metrics and types of EEG biomarkers, simulated dose effects were linear, in line with the gradual α5-PAM dose-effects seen behaviourally in rodents (Prevot et al. 2019). In comparison, the simulated non-selective PAM did not recover the power spectral profile. This is in agreement with previous studies showing that non-selective benzodiazepines and benzodiazepines selective for α1 subunits have been associated instead with increases in EEG delta and beta rhythms in humans during resting-state (Jobert, Schulz, and Jähnig 1995; Jobert and Wilson 2015; Premoli et al. 2017), and with a slowing of the peak frequency over time (Jobert et al. 1995). These findings are consistent with the poor effectiveness of non-selective PAM at treating depression, as suggested by previous clinical studies (Parker and Graham 2015).

We constrained the models with the effect of 3 μM of the α5-PAM compound, G-II-73, which was in the previously demonstrated range for selectively targeting α5 subunit receptors compared to higher concentrations that more strongly target α1, α2, and α3 subunits (Bernardo et al. 2022; Prevot et al. 2019). We assumed that the α5-PAM effect on tonic inhibition conductance, which we estimated from the human neuronal recordings, would also occur with synaptic conductance of SST→Pyr connections because of the localization of α5 subunits in those synapses (Ali and Thomson 2008; Davenport et al. 2021; Donato et al. 2023; Schulz et al. 2018). The proportion of α5-PAM effects on synaptic or tonic inhibition would depend on trafficking of α5 subunits to extrasynaptic and synaptic locations, which is highly dynamic and activity-dependent (Davenport et al. 2021; Jacob 2019). Further data of α5-PAM effect on synaptic vs tonic inhibition could thus refine the models and increase the accuracy of their predictions.

We tested α5-PAM effect on cortical function using signal detection metrics that relate to deficits in depression (Huang, Thompson, and Paulus 2017; Koetsier et al. 2002; Tsourtos, Thompson, and Stough 2002), possibly due to increased noise in cortical processing as a result of reduced SST interneuron inhibition (Yao et al. 2022). Administration of α5-PAM and characterization of its effects on cognition has previously only been done in rodents (Koh et al. 2013; Prevot et al. 2019). Whereas both sensory and memory impairments are thought to be linked to increased noise in cortical processing, although in different corresponding cortical regions (Northoff and Sibille 2014; Prevot and Sibille 2021), future simulation studies will benefit from testing α5-PAM effects on other, more complex, cognitive functions than those we have modeled.

As in previous studies, we used models of human cortical L2/3 microcircuits to study effects on cortical microcircuit function (Yao et al. 2022) and EEG signal (Mazza et al. 2023). These serve as prototypical models of human cortical microcircuitry, as supported by their ability to reproduce key aspects of human resting state EEG recordings (Mazza et al. 2023), due to the use of realistic human neuronal morphologies, human synaptic properties, and the key neuron types in the microcircuit. Further support for the predictive power of the L2/3 microcircuits stems from L2/3 being closest to the EEG electrode and thus a major contributor to EEG signals (Buzsáki, Anastassiou, and Koch 2012). Future expanded models that include layers 4 and 5 could help refine our estimated EEG biomarkers of α5-PAM effects, by including the additional complexities of interlaminar oscillatory communication (Florez et al. 2015) as well as oscillatory dynamics present in deeper cortical layers (Guet-McCreight et al. 2022; Roopun et al. 2006). Although we simulated L2/3 microcircuits ∼6-7 times smaller in terms of cell numbers than the true size (for computational efficiency), the down-sampling involved proportional changes in both excitatory and inhibitory neurons, which maintained the overall excitatory–inhibitory balance of the network. We do, however, expect our biomarkers will largely hold, so that any refinements will rather serve to provide additional biomarker candidates such as phase-amplitude coupling between different frequency bands originating from different layers (Florez et al. 2015). Whereas we used models of depression microcircuits with reduced SST interneuron inhibition that were constrained with expression data from depression patients (Yao et al. 2022), other mechanisms of depression include morphological atrophy and reduced spine density (Banasr, Dwyer, and Duman 2011; Bernardo et al. 2022). However, we note that chronic α5-PAM exposure was shown to be effective in directly recovering these morphological features (Bernardo et al. 2022; Prevot et al. 2021), therefore the translation of α5-PAM effects on these depression mechanisms to humans would be more trivial.

Altogether, our results provide the first demonstration that α5-PAM intervention in the context of human depression has the potential at reversing altered SST-mediated neurotransmission and alleviate related cognitive impairments. Our study also presents a first in-silico testing of pharmacology systematically on detailed models of human cortical microcircuits, which we hope will also be of service in future efforts.

## Methods

### Electrophysiology data

We used whole-cell voltage-clamp recordings of tonic inhibition current in the presence of GABA only, and in the presence of α5-PAM + GABA, in human cortical L2/3 Pyr neurons (10 cells: 9 cells from 3 male subjects, 1 cell from 1 female subject) from patients undergoing a selective amygdalohippocampectomy (Lozano, Gildenberg, and Tasker 2009). As described in previous work (Howard et al. 2022), the resected cortical tissue was considered healthy as it was located outside of the site of epileptogenesis. Written informed consent was obtained from all participants, in accordance with the Declaration of Helsinki and the University Health Network Research Ethics board.

The data was collected using surgery resection, solutions, tissue preparation, and recording equipment described previously (Chameh et al. 2021; Howard et al. 2022; Yao et al. 2022). Neocortical tissue resected during anterior temporal lobectomy was immediately submerged in ice-cold (∼4°C) cutting solution and transferred to a recording chamber within 20 minutes. After sectioning the tissue, the slices were incubated for 30 min at 34 °C in standard artificial cerebrospinal fluid (aCSF) (in mM): NaCl 123, KCl 4, CaCl_2_.2H_2_O 1.5, MgSO_4_.7H_2_O 1.3, NaHCO_3_ 26, NaH_2_PO_4_.H_2_O 1.2, and D-glucose 10, pH 7.40 and bubbled with carbogen gas (95% O_2_-5% CO_2_) and had an osmolarity of 300-305 mOsm.

For recordings, slices were transferred to a recording chamber mounted on a fixed-stage upright microscope (Axioskop 2 FS MOT; Carl Zeiss, Germany). Slices were continually perfused at 4 ml/min with standard aCSF at 32-34 °C. Whole-cell patch-clamp recordings were obtained using a Multiclamp 700 A amplifier and pClamp 10.6 data acquisition software (Axon instruments, Molecular Devices, USA). Subsequently, electrical signals were digitized at 20 kHz using a 14140A digitizer. For voltage-clamp recordings of tonic current, low-resistance patch pipettes (2– 4 MΩ) were filled with a CsCl-based solution containing (in mM) 140 CsCl, 10 EGTA, 10 Hepes, 2 MgCl_2_, 2 Na_2_ATP, 0.3 GTP, and 5 QX314 adjusted to pH 7.3 with CsOH. The junction potential was calculated to be 4.3 mV and the holding potential was -74.3 mV after junction potential correction. As in previous studies (Asgari et al. 2016; Schulz et al. 2018), in this configuration, 5 μM GABA, 25 μM AP5, 10 μM CNQX, and 10 μM CGP-35348 were first applied to generate larger GABA-dependent currents and assess tonic inhibition currents while also blocking AMPA, NMDA, and GABA_B_ mediated currents. 3 μM of α5-PAM compound GL-II-73 (Prevot et al. 2019) was then applied to assess tonic inhibition current in the presence of α5-PAM, followed by 50 μM of picrotoxin to block GABA_A_ mediated currents and assess endogenous current output during voltage-clamp recordings without any synaptic activity.

### Human cortical microcircuit models in health and depression

We used morphologically- and biophysically-detailed models of human L2/3 cortical microcircuits in health and depression described previously (Yao et al. 2022). Briefly, these microcircuit models were comprised of 1000 neurons (80% Pyr, 5% SST, 7% PV, and 8% VIP) distributed across a 500×500×950 µm^3^ volume and simulated using NEURON (Carnevale and Hines 2006) and LFPy (Hagen et al. 2018). The neuronal morphology reconstructions of the multi-compartment models were obtained from the Allen Cell Types database (Gouwens et al. 2018), and the models were fitted using multi-objective optimization (Hay et al. 2011; Van Geit et al. 2016) with either single cell data from the Allen Brain Institute (putative PV, SST, and VIP inhibitory neuron fits; Gouwens et al., 2018) or population Pyr neuron data from the Krembil Brain Institute (Chameh et al. 2021; Howard et al. 2022). Synaptic parameters in these models were fit to human data where possible (Komlósi et al. 2012; Obermayer et al. 2018; Seeman et al. 2018; Szegedi et al. 2016) and to curated rodent data otherwise (Ramaswamy et al. 2015). In terms of connections onto Pyr neurons which had two types of dendritic trees (basal and apical), the Pyr→Pyr excitatory synapses were placed on both basal and apical dendritic compartments, the PV→Pyr inhibitory connections were placed on basal dendritic compartments, the SST→Pyr inhibitory connections were placed on apical dendritic compartments. Depression microcircuits were modelled by reducing the conductance of SST interneuron synaptic and tonic inhibition onto all cell types by 40% (Yao et al. 2022). For Pyr neurons in the depression model, tonic inhibition conductance was reduced by 40% on only apical dendritic compartments. For each interneuron type in the depression model, the contributions of SST interneurons to tonic inhibition were estimated and this contribution was reduced by 40%. Randomizing the circuit comprised of sampling synaptic connections, neuron positions in space, background noise input, and spike timing of thalamic inputs (see section below). Full model details are available in Yao et al (2022), and the models are openly accessible online: https://doi.org/10.5281/zenodo.5771000.

### Modelling microcircuit baseline and response activity

As in previous work (Yao et al. 2022), the microcircuit generated spike rates at baseline and during response in line with the different neuron types in vivo (Gentet et al. 2012; Teleńczuk et al. 2017; Yu et al. 2019), and each neuron received random background excitatory inputs using Ornstein-Uhlenbeck (OU) point processes (Destexhe et al. 2001).

As in Yao et al. (2022), we reproduced response rates using excitatory AMPA/NMDA synapses with the same synaptic dynamics and number of contacts as the cortical excitatory synapses. 55 Pyr neurons were stimulated in the basal dendrites, with 2 – 4 ms delay post-stimulus and a conductance of 4 nS. 35 PV interneurons were stimulated with a delay of 2 - 2.5 ms and a conductance of 2 nS. VIP interneurons were stimulated in two groups and phases: early (65 VIP interneurons, delay = 0.5 - 4.5 ms, conductance = 2.8 nS) and late (80 VIP interneurons, delay = 7 - 12 ms, conductance = 2.2 nS). Average post-stimulus rates were calculated over the 5 - 55 ms window after stimulus onset.

### Tonic inhibition models

As in Yao et al (2022), we used a model for outwardly rectifying tonic inhibition (Bryson et al. 2020) as well as the tonic inhibition conductance values that had previously been fitted to capture the current magnitudes recorded in human L2/3 Pyr neurons in the presence of GABA only (see electrophysiology methods above and Yao et al., 2022). We simulated the experimental conditions by setting the inhibitory chloride reversal potential to -5 mV (consistent with the experimental solutions), setting the holding potential to -75 mV in voltage-clamp mode, and tuning the tonic inhibition conductance on all Pyr neuron somatic and dendritic compartments to reproduce the target experimental tonic inhibition current amplitude (resulting in *G_tonic_* = 0.938 mS/cm^2^). The same *G_tonic_* value was used for the interneurons since the total tonic inhibition current recorded in interneurons is similar to that of Pyr neurons after correcting for cell capacitance (Scimemi et al. 2006).

### α5-PAM models

We estimated the *G_tonic_* modulation on Pyr neuron apical dendrites during application of α5-PAM (resulting in *G_tonic_* = 1.498 mS/cm^2^) using the experimental tonic inhibition currents as target magnitudes, and the simulation settings as described for tonic inhibition models above. All simulated current magnitudes were calculated relative to the endogenous current magnitude generated in the condition where *G_tonic_* is set to 0 mS/cm^2^, corresponding to the picrotoxin condition in the experimental methodology. We then applied the estimated α5-PAM modulation effect on both tonic and synaptic inhibition, corresponding to *G_tonic_* and SST→Pyr synaptic conductance (*GSST→Pyr*) in the microcircuit models. We simulated different doses of α5-PAM ranging from 25% to 150% of the estimated modulation effect of the experimental reference dose.

### Non-selective PAM models

We modeled the action of non-selective GABA_A_ receptor PAM (i.e. benzodiazepines broadly non-selective for α1-, α2-, α3- and α5 subunit-containing GABA_A_ receptors) by applying the same magnitude of estimated α5-PAM modulation, but to all synaptic and tonic inhibition connections in the microcircuit.

### Failed/false signal detection rates

We computed error rates in stimulus processing with our microcircuit models as in previous work (Yao et al. 2022), by first fitting the pre-stimulus firing rate distributions (computed using a 50 ms sliding window, sliding in 1 ms intervals, over a 3s pre-stimulus period) to skewed normal distributions for each of the 200 randomized microcircuits (n = 2,951 windows × 200 microcircuits pre-stimulus). We then fitted the post-stimulus firing rate distribution (in the 5-55 ms period post-stimulus) across all the 200 randomized microcircuits (n = 200 windows). The intersection point between pre-stimulus distribution and the post-stimulus distribution was chosen to be the stimulus detection threshold, in line with optimal decision theory (Berger 1985). Probability of false detections was computed as the integral of the pre-stimulus distribution above the detection threshold divided by the integral of the entire pre-stimulus distribution. The probability of failed detections was computed as the integral of the post-stimulus distribution under the detection threshold divided by the integral of the entire post-stimulus distribution.

### Simulated microcircuit EEG and power spectral analyses

We simulated dipole moments and corresponding EEG time series data generated by our microcircuit models (25s duration per simulation) using the same methodologies as in previous work (Hagen et al. 2018; Mazza et al. 2023). Specifically, we used a four-sphere volume conductor model (representing grey matter, cerebrospinal fluid, skull, and scalp with radii of 79 mm, 80 mm, 85 mm, and 90 mm, respectively) that assumed homogeneous, isotropic, and linear (frequency-independent) conductivity. The conductivity for each sphere was 0.047 S/m, 1.71 S/m, 0.02 S/m, and 0.41 S/m, respectively (Mazza et al. 2023; McCann, Pisano, and Beltrachini 2019). We computed EEG power spectral density using Welch’s method (Welch 1967) from the SciPy python module, and quantified power spectral features by computing the integral of the power spectral densities for theta (4 - 8 Hz), alpha (8 - 12 Hz), and lower beta (12 - 21 Hz) range frequencies.

As in previous work (Mazza et al. 2023), we decomposed the EEG power spectral densities (in the 2 - 30 Hz range) into aperiodic and periodic components using an algorithmic parameterization method (Donoghue et al. 2020). The aperiodic component was a 1/f function parameterized by vertical offset and exponent parameters. As an additional quantification metric, we computed the integral of the broadband (3 - 30 Hz) frequency range in the aperiodic component (or area-under the curve, AUC). Overlying the aperiodic component, we fitted the periodic oscillatory component with up to 4 Gaussian peaks which were defined by center frequency, bandwidth (min: 2 Hz, max: 12 Hz), and power magnitude (relative peak threshold: 2, minimum peak height: 0.5). We quantified periodic features by computing the integral of the periodic component for theta, alpha and lower beta range frequencies.

We also computed the spiking power spectral density of Pyr neurons by converting the spike times into binary spike train vectors, which we then summed across all Pyr neurons. Power spectral density was then computed from the summed spike train vectors using Welch’s method (Guet-McCreight and Skinner 2019; Welch 1967; Yao et al. 2022). For both Pyr neuron spiking and EEG, we computed power spectral density with nperseg = 140,000 sampling points, which was equivalent to 3.5s time windows. For spiking and EEG power spectral density vectors, as well as the aperiodic and periodic vectors, across random seeds we computed the bootstrapped (100 iterations) means and 95% confidence intervals for each frequency.

## Acknowledgements

AGM and EH thank the Krembil Foundation for their generous funding support. AGM thanks the Yuet Ngor Wong Awards for funding support. TAV also thanks the generous support from CAMH Discovery Fund and Kavli Foundation. As well, we are immensely grateful to our neurosurgical patients and their families for consenting to the use of their tissue samples for research. Special thanks to A. Sherrington for contributing the drawing in Fig. 4A.

